# Detecting consistent patterns of directional adaptation using differential selection codon models

**DOI:** 10.1101/072405

**Authors:** Sahar Parto, Nicolas Lartillot

## Abstract

**Background:** Phylogenetic codon models are often used to characterize the selective regimes acting on protein coding sequences. Recent methodological developments have led to models explicitly accounting for the interplay between mutation and selection, by explicitly modelling the amino acid fitness landscape along the sequence. However, thus far, most of these models have assumed that the fitness landscape is constant over time. Fluctuations of the fitness landscape may often be random or depend on complex and unknown factors. However, some organisms may be subject to systematic changes in selective pressure, resulting in reproducible molecular adaptations across independent lineages subject to similar conditions.

**Results:** Here, we developed a codon-based differential selection model, which aims to detect and quantify the fine-grained consistent patterns of adaptation at the protein-coding level, as a function of external conditions experienced by the organism under investigation. The model parameterizes the global mutational pressure, as well as the site- and condition-specific amino acid selective preferences. This phylogenetic model is implemented in a Bayesian MCMC framework. After validation with simulations, we applied our method to a dataset of HIV sequences from patients with known HLA genetic background. Our differential selection model detects and characterizes differentially selected coding positions specifically associated with two different HLA alleles.

**Conclusion:** our differential selection model is able to identify consistent molecular adaptations as a function of repeated changes in the environment of the organism. These models can be applied to many other problems, ranging from viral adaptation to evolution of life-history strategies in plants or animals.

## Background

Statistical models of molecular evolutionary processes are now widely used to analyze the interplay between mutation and selection. Often, these models are formulated at the codon level, thus relying on the contrast between synonymous and non-synonymous substitutions to leverage out an estimation of the strength of selection acting at various levels (nucleotide, amino acids, codon usage) of protein coding sequences. The first codon models, proposed independently by Goldman and Yang [1] and Muse and Gaut [2], relied on a simple aggregate parameter, ω=*dN/dS*, to capture the overall strength of selection, globally over the protein coding sequence and over the phylogenetic trees. Subsequent elaborations on these original models allowed for variation in *dN/dS* among sites [3, 4] or among lineages [5], thus increasing the sensitivity and the resolution of the detection of selective regimes. However, all of these models still do not discriminate between alternative amino acids. Instead, they essentially put all non-synonymous substitutions on the same level [6].

In this direction, Halpern and Bruno [7] and also Thorne et al [8] have proposed an alternative codon modelling strategy, allowing for site- and amino acid-specific selective effects. Their model also has a clear mechanistic interpretation, being derived from first principles of population genetics. Specifically, in their model, the rate of substitution between codons is seen as the product of the mutation rate and the fixation probability. In turn, the fixation probability is made explicitly dependent on the selection coefficient of the mutation under consideration. Selection coefficients are obtained from an explicit fitness landscape, in which the fitness of each amino acid is allowed to be different at each coding site. Technically, the model therefore invokes, at each coding site, a normalized vector of 20 amino acid fitness coefficients, collectively referred to as the site-specific fitness profile. In the original version of Halpern and Bruno, site-specific amino acid fitness profiles were empirically estimated based on observed amino acid frequencies. Since then, a statistically more sophisticated version of this model was developed in a Bayesian framework by Rodrigue et al [6], using a non-parametric approach to integrate over the uncertainty about site-specific selective features (now seen as random-effects across sites), and to capture the unknown law of amino acid fitness profiles across sites. The importance of accounting for modulation of selection across sites by introducing site-specific amino acid fitness profiles was demonstrated by Bayes factor computation and posterior-predictive tests [6]. Of note, more phenomenological variants of this modeling approach, also with site-specific amino acid fitness contributions but without the population-genetic justification of Halpern and Bruno’s paradigm, have been explored [6-9].

This modeling approach, although fairly complex, still leaves an important aspect of protein evolution aside, by assuming that the fitness landscape is constant through time. Yet, many ecological situations clearly suggest that fitness landscapes undergo important fluctuations through time [10]. Two alternative approaches are possible, to relax this specific assumption. First, fluctuations of the fitness landscape could be modelled as a purely latent effect (e.g. Markov-modulated models) [11], thus without relying on any extra information about the environmental or ecological drivers of the fluctuations. Secondly, in some situations, empirical knowledge is available, in terms of varying conditions across sampled genetic sequences. In this context, it is, in principle, possible to explicitly model condition-specific amino acid fitness modulations. The present work is an attempt at modeling such effects.

A clear-cut example where robust empirical knowledge about varying selective environments is available is the evolution of viral sequences as a function of the genetic background represented by the hosts. For example, the analysis of patterns of selection, using *dN/dS* codon models in a phylogenetic maximum likelihood framework, has shown the substantial role of fluctuating selection in the emergence of new mutations and the ability of HIV-1 to escape from immune system [12-14]. HIV-1 is capable of evading the CTL (Cytotoxic T-Lymphocyte) response because of its rapid rate of mutation in HLA-restricted epitopes, called escape mutation. Escape mutation gives the virus the ability to adapt under different selective forces in different individuals and in response to drugs, which makes the design of a vaccine very difficult. Therefore, understanding the evolution of HIV-1 within the human body, which is both rapid and under strong selection, helps designing more effective vaccines against HIV-1 and control its evolution. On the other hand, the high rate of mutation of HIV-1 enables the virus to produce genetically distinguished population in each host, called quasispecies [15], which let the evolutionary studies possible within the HIV-1 population. The correlation between HLA alleles and HIV polymorphisms has been paid a lot of attention in recent years, from population-based studies [16-18] to studies taking phylogeny into account [19, 20]. The Phylogeny Dependency Network study accounts for phylogeny, codon correlation and HLA linkage disequilibrium to analyze HLA-mediated escape in HIV-1 [21]. However, this method only takes the information of the tips of the phylogenetic tree into account. More fundamentally, it does not rely on an explicit model of the underlying molecular evolutionary processes. Another phylogenetic model has been used by Tamuri et al [22] to identify host dependent selective constraints for flu viruses. These authors specified different host-dependent substitution rates along the phylogenetic tree, and used a maximum likelihood approach, combined with a likelihood-ratio test, to identify positions under differential selection between hosts. One potential short-coming of the modeling approach used in [22]] is that it is formulated directly at the amino acid level. Therefore, a more explicit codon modeling approach could be used as an alternative, to tease out in a more principled manner, the respective contributions of mutation and selection processes in the observed patterns of sequence evolution.

In this direction, we now introduce a codon model able to capture site- and condition-specific amino acid fitness effects. In this Bayesian model, which we call differential selection (DS) model, a site and branch heterogeneous selection factor is invoked to estimate the substitution rate at the codon level of aligned HIV-1 sequence. As the population-genetics of viral populations is complex and difficult to model quantitatively, we explored two alternative strategies for deriving the codon substitution process, either using a phenomenological approach, or using a mechanistic derivation as in Halpern and Bruno. Our differential selection model was then used to investigate how the fluctuating environment provided by the diversity of human HLA background affects HIV-1 sequence evolution. We illustrate how our approach finds consistent patterns of viral adaptation, in terms of how selection acts at specific positions, modulating amino acid preference as a function of the HLA background.

## Materials and Methods

### HIV-1 data

333 HIV-1 DNA sequences of subtype B from Gag region of HIV-1 from 41 patients with known HLA types were obtained from Los Alamos National Laboratory (LANL) HIV-1 sequence database [23]. Each patient has on average 8 sequences. Information about the HLA types of the patients was also downloaded. About 35% of the sequences are from HLA B57+ patients. Recombinant sequences were excluded from the study (they were removed by the software in the download process). The amino acid alignment of the sequences provided by the source was downloaded, manually corrected (misplaced amino acids were relocated and misaligned regions were deleted) and used for back aligning the DNA sequences at the codon level.

### Phylogenetic tree estimation

Primarily for computational reasons, the method introduced here assumes a fixed tree topology. However, owing to the relatively short length of the coding sequences that were used, this topology may not be known with high confidence. In addition, there is the question of whether the sequences corresponding to a given patient should form a monophyletic group. This may not be the case because of tree reconstruction errors, a problem which can be alleviated simply by constraining the monophyly of each patient during the tree reconstruction. However, non-monophyly could be real, being caused by complicated infection patterns between individuals. In this case, constraining the monophyly might introduce mis-specifications in the reconstructed tree topology.

To check the robustness of our method to these potential sources of error, we tested alternative methods for reconstructing the phylogenetic tree and conducted independent analyses under these alternative tree topologies. Specifically, a first tree topology (T1) was obtained directly from the LANL website. This tree was estimated using the neighbor joining algorithm [24]. A second tree (T2) was reconstructed using MrBayes [25, 26], under the GTR+Gamma substitution model and constraining the monophyly of the groups corresponding to sequences belonging to a given patient. A third tree (T3) was estimated, still using MrBayes, under the same substitution model, but without imposing any constraint on the tree topology. In MrBayes, we ran MCMC chain for 1500000 cycles (the average standard deviation of split frequencies reaches the value less than 0.05, and the Potential Scale Reduction Factor (PSRF) [27], which should approach 1.0 as the two runs converge, was equal to 1.001 and 1.000 for the two analyses).

In the case of tree T1 and T3, we observed 20 and 23 cases of non-monophyletic patients, respectively. In both cases, we applied a greedy algorithm for excluding the smallest possible set of sequences such that each patient is then represented by a monophyletic group of sequences. This was done using the following recursive procedure: first, the number of sequences from each host pending from (downstream to) each node was determined recursively at each node, from the tips toward the root. During this recursive scan, wherever a group pending from a given node was not monophyletic, the sequences belonging to the host with the smallest number of sequences pending from that node were flagged. Finally, in a backward recursive scan of the tree, from root to tips, the flagged sequences were removed from the dataset. Application of this method leads to the elimination of 20 and 23 out of 333 sequences in the cases of tree T1 and T3. Lastly, for the three topologies, the branches of the phylogenetic tree were divided into 4 conditions according to the host HLA types (see below).

### Model

#### Notations –

We consider a coding sequence of length *N* (N being the number of coding positions, or equivalently *3N* is number of nucleotide sites). The number of conditions (e.g. HLA types) is defined by *K*. All the indices used in this paper conform to the following conventions:

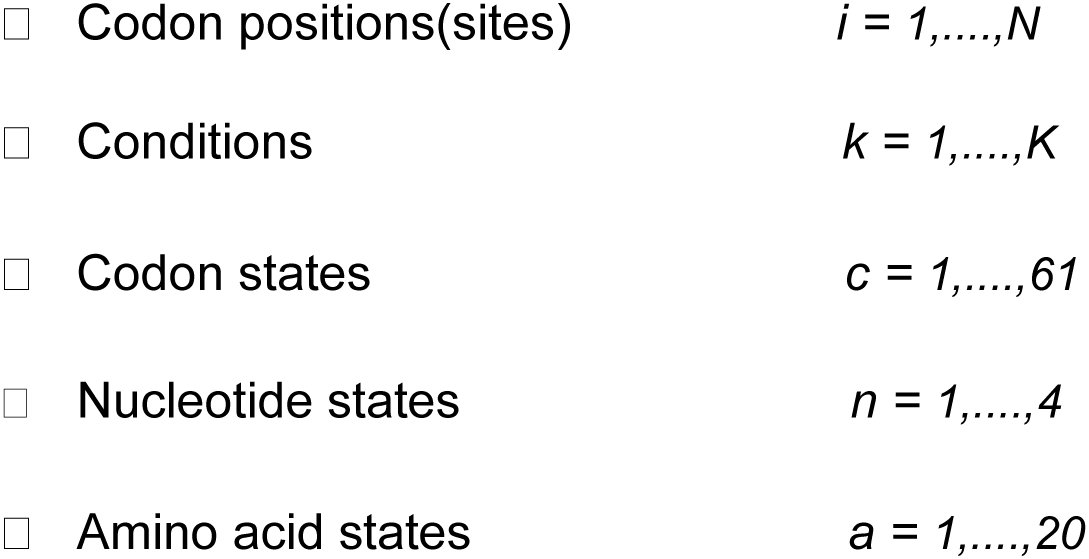

#### Model of codon substitution –

The rate of evolution by point substitution is the result of a complex interplay between mutation, selection and random drift. Drawing inspiration from previous developments in statistical molecular evolution [1, 2, 6, 7, 9], we modelled this complicated process at the codon level, as a multiplicative combination of mutation rates and selective effects (the latter implicitly including the contribution from random drift).

The mutation process is assumed to be homogenous over time and along the sequence. It is modelled as a Markovian general time-reversible process, parameterized in terms of the relative exchange rates between nucleotides (*ρ*) and the stationary probability (equilibrium frequency) of the target nucleotide (*π*). Thus, the rate of substitution from nucleotide *n*_1_ to nucleotide *n*_2_ is equal to:

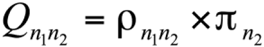

The set of relative exchangeabilities between nucleotides is constrained to be symmetric:

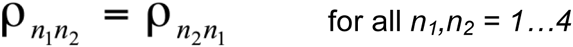

In addition, it is normalized:

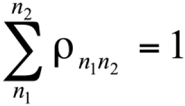

The vector *π* of equilibrium frequencies is also normalized:

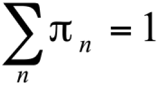

The selective forces, on the other hand, are both condition- and position-specific. The modulations across conditions and positions are mediated exclusively by the encoded amino acid sequence. Accordingly, for each position *i* and each condition *k*, we introduce an array of 20 non-negative fitness factors 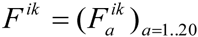, one for each amino acid. In the following, these 20-dimensional vectors will be referred to as amino acid *fitness profiles*. Thus, we have distinct fitness profiles across positions, and for a given position, the fitness profile over the 20 amino acids is further modulated across conditions. How these fitness profiles are defined in practice is explained in more detail below (section; Definition of the amino acid selective effects).

Given a mutation matrix and a set of amino acid fitness profiles, we considered two alternative approaches for expressing substitution rates between codons as a function of the fitness of the amino acids. The first is a phenomenological approach, while the second is more mechanistic in its inspiration.

#### Phenomenological model (M1) —

the phenomenological model is similar, in its general form, to the models explored by Rodrigue et al [6], or, in a slightly different parameterization, to the models considered in Robinson et al [9]. Specifically, consider a given position *i* along the sequence, and a given condition *k* along the tree. Consider also two codons, *c*_1_ and *c*_2_, differing only at one position and with nucleotides *n*_1_ and *n*_2_ at that position. These two codons encode for amino acids *a*_1_ to *a*_2_, respectively. Then, the rate of substitution between these two codons is given by:

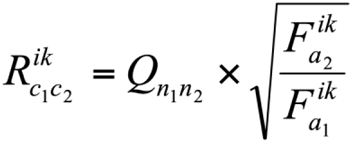

Thus, according to this model, the rate of substitution is proportional to mutation rate, while being influenced by the selection operating at the amino acid level, through the fitness factors 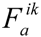: the substitution rate is higher (resp. lower) than the neutral substitution rate if the fitness of the final amino acid is greater (resp. smaller) than the fitness of the initial amino acid. Note that, if the two codons are synonymous, i.e. if *a*_1_=*a*_2_, then the substitution rate is simply equal to the mutation rate defined by the nucleotide transition matrix *Q*. Finally, the model considers only point substitutions, and therefore, the substitution rate is assumed to be equal to zero between codons differing at more than one nucleotide position. Thus, altogether:

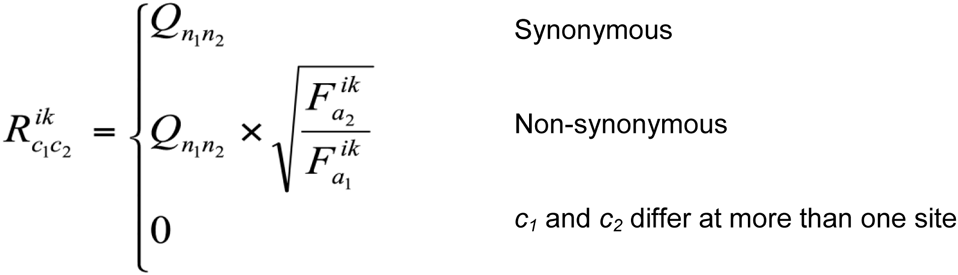

#### Mechanistic model (M2) —

The second approach is inspired by a mechanistic argument based on first principles of population genetics, as initially suggested by Halpern and Bruno [7]. Suppose again the substitution rate between codon *c*_1_ to *c*_2_ at site *i* and condition *k*. First, we define a scaled selection coefficient, associated with codon *c*_2_, seen as a mutant in the context of a population in which the wild-type allele is *c*_1_. This scaled selection coefficient is given by:

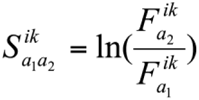

Then, the rate of substitution between codon *c*_1_ and *c*_2_ is given by the product of the mutation rate and the relative fixation probability *P* (i.e. relative to neutral). This fixation probability is itself dependent on the scaled selection coefficient. Using the classical diffusion approximation, this relative fixation probability can be expressed as:

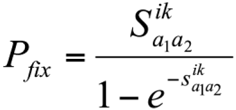

So that the rate of substitution between codons is given by

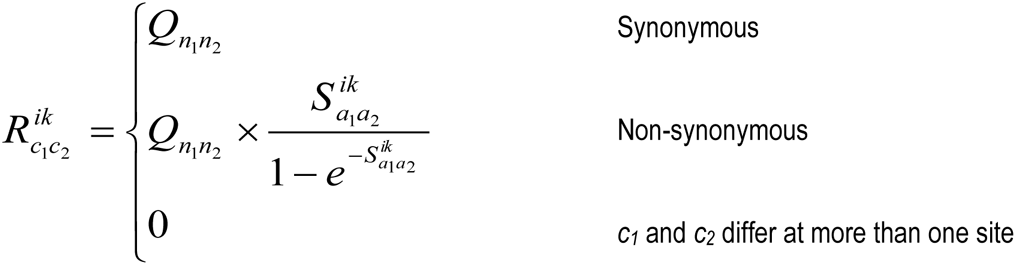

Again, we see that the rate of substitution is higher (resp. lower) than the neutral substitution rate if the non-synonymous mutation leads to an increase (resp. a decrease) in the fitness of the sequence.

#### Definition of the amino acid selective effects —

In principle, the amino acid fitness profiles associated to each site and each condition, 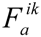, could be considered as independent arrays, both across sites and across conditions. However, most of the amino acid conservations (due to purifying selection) observed along the sequence is in fact condition-independent. Against this globally invariable fitness background, the modulations of the fitness landscape induced by condition-dependent effects (such as the HLA type of the host) are likely to be comparatively small. In this context, considering amino acid selective effects as totally independent parameters across conditions would imply that the invariable background would be re-estimated independently for each condition, potentially resulting in a loss of statistical power. Therefore, as a more powerful alternative, we explicitly defined amino acid selection in terms of a log-additive superposition of a global background and condition-dependent differential selective effects, as follows. First, a baseline or global fitness profile is defined for each position. That is, for position *i*, we define a 20-dimensional vector 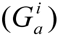 for *a=1…20*. This vector is drawn from a uniform Dirichlet distribution independently at each site. This baseline defines the fitness landscape under condition 0, which is therefore taken as our reference condition (black branches in Figure 1).

**Figure 1.**
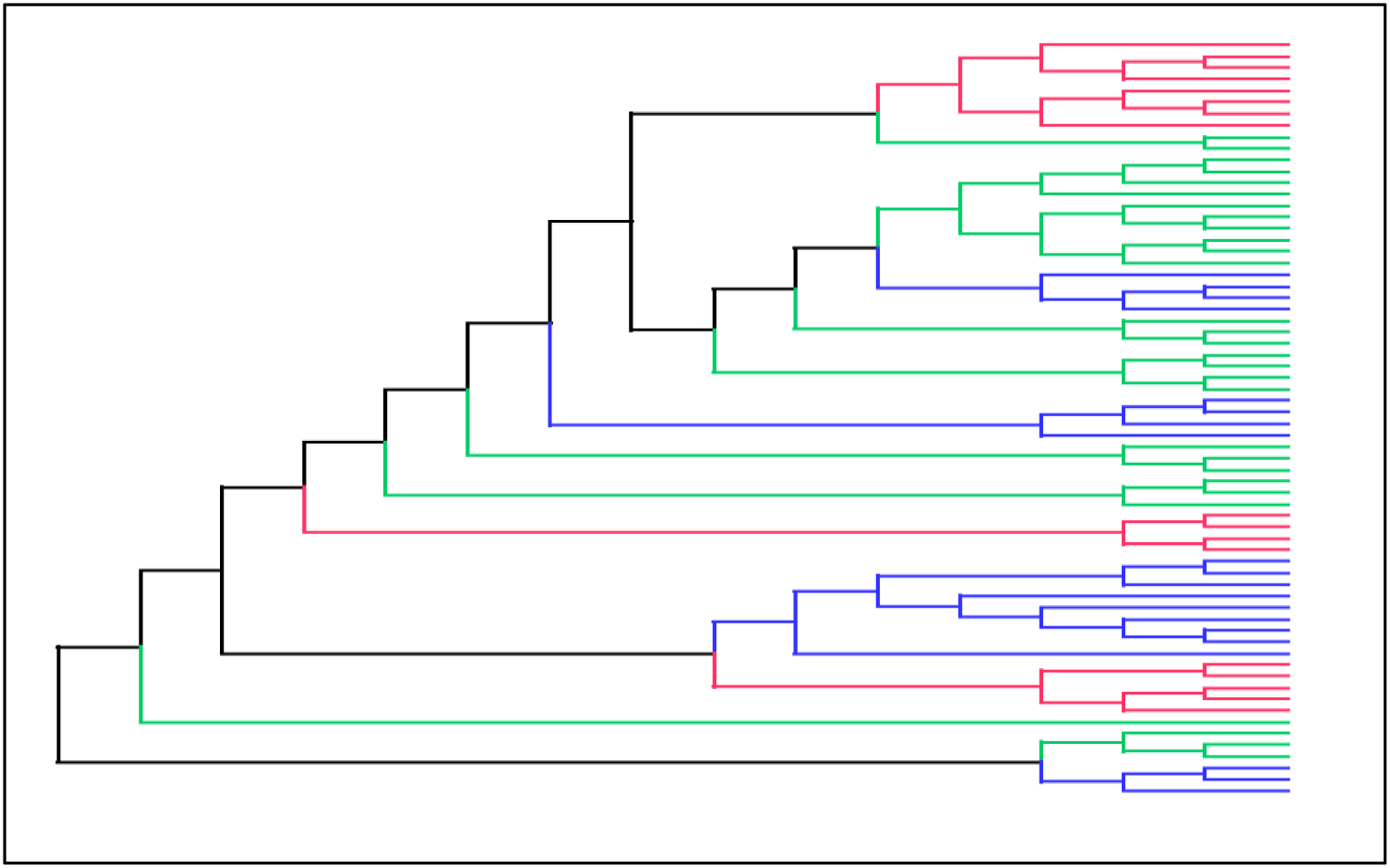
Illustrative phylogenetic tree of 313 HIV-1 Gag sequences. Different colors along the tree show different selection regimes for the corresponding sequences. Black for between-patient, green for within-patient, red and blue for HLA B57 and HLA B35 dependent categories, respectively.

Next, selection is modulated across conditions through the use of condition-specific differential selection profiles. Thus, for position *i* in condition *k*, we define a 20-dimensional vector 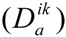, for *a=1…20*. Unlike the baseline profiles, which are positive (and sum to 1), those differential selection effects can be positive or negative. A positive (resp. negative) coefficient means that the fitness of the corresponding amino acid is increased (resp. decreased) in the target condition, compared to the reference condition. The differential selection profiles are drawn *iid* from a Normal distribution of mean 0 and condition-specific variance 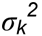.

Altogether, the condition-specific fitness profiles are constructed as follows:

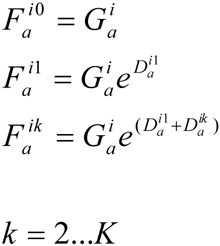

Note that we have used a two-level system for introducing the differential effects (i.e. a different equation for *k=0* and *k>0*). This is motivated by the fact that we need to discriminate both among branches that are between hosts and within the same host, and among hosts with differing HLA backgrounds. Thus, it reflects the differential between within-host (*D*^*i*1^) and between-host (*G^i^*) selection regions, while representing specific selective features more specifically associated to differing HLA backgrounds (*D*^*ik*^)_*k*=2…*K*_. In the case of HIV-1, we consider 2 focal HLA backgrounds (B57+ and B35+), against a default B57−/B35− background. Thus, we define a total of 4 different conditions (*K=4*), and the branches of the tree are partitioned according to 4 different selection regimes: between hosts (*k=0*), within B57−/B35− hosts (*k=1*), within B57+ (*k=2*) and B35+ (*k=3*) patients (Figure 1).

An important point should be emphasized concerning the statistical formalization of the fitness landscape and of its modulations across sites and across conditions. Conceptually, the arrays of global and condition-specific fitness effects should be considered, not as parameters, but as random-effects across sites, which are integrated over a distribution (respectively, a Dirichlet and a normal distribution for the global and differential effects). This integration is done implicitly, through the MCMC sampling (see below). As a result, the aim of the model introduced here is not to achieve accurate and asymptotically consistent point estimation of site- and condition-specific fitness effects: in most cases, the information for inferring such fitness effects will be limited. Instead, it is to draw inference based on the complete posterior distribution. A more specific objective is to single out those relatively few cases for which there are sufficient information to infer, with high posterior probability, the presence of a differential selective effect between two conditions. One important desirable property of this type of inference is to allow for a reasonably good control of the fraction of false discoveries among those cases that are selected based on a high posterior probability of a differential effect. This is something which is investigated through posterior predictive simulations.

#### Priors-

The topology (*τ*) of the tree is fixed. The parameters of the model consist of branch lengths, *l_j_* (1<*j*<2*N*-3 where *N* is the number of sequences), nucleotide exchangeabilities, *ρ* and nucleotide equilibrium frequencies, *π*. The priors that we used are as follows: on branch lengths: a product of independent Exponentials of mean *λ*; the hyperparameter *λ* is from Exponential distribution of mean 0.1; on relative exchangeability rate: a product of Exponentials of mean 1; on mutational equilibrium frequency: a uniform Dirichlet distribution. As mentioned above, the site-specific fitness profiles (*G*) and differential fitness effects (*D*) are random-effects, integrated over Dirichlet and normal distributions, respectively.

#### MCMC-

We used Markov Chain Mont Carlo (MCMC) to sample the parameters of the model from their joint posterior distribution. We used a graphical model environment previously introduced in [28], heavily relying on data augmentation and parameter expansions methods, such as described in particular in [29]. Briefly, a MCMC cycle consists of an alternation between two steps: first, a detailed substitution history at each coding site is Gibbs-sampled, from the posterior distribution conditional on the current parameter configuration. Second, conditional on these augmented data, the parameters and the random-effects across sites are updated through a large series of Metropolis-Hastings moves, cycling over all parameters or random variables of the model.

For nucleotide equilibrium frequencies *π* and global fitness profiles *G,* which are under the constraint that they should sum to 1, we used constrained move as explained in [28]. For the branch lengths *l* and the exchangeabilities *ρ*, which are positive real numbers, the multiplicative moves were used for their updates [28]. After 500 points of burn-in are removed, posterior estimates are estimated by averaging over the remaining of the MCMC chain (approximately 1500 points).

### Simulation analyses

Simulations were conducted using the posterior predictive formalism, as described in [30, 31], using the HIV dataset as a template, and under two versions of the model: (1) with only one condition across the whole tree (thus representing the null hypothesis of no differential effect across conditions); and (2) with the 4 conditions described above (between and within patients, with differing HLA backgrounds). In both cases, the phenomenological (M1) and the mechanistic (M2) models were investigated. In each of these four cases, two independent runs of the MCMC were conducted on the empirical HIV dataset. Then posterior predictive simulations were conducted on 5 parameter configurations sampled from the posterior distribution (5 points regularly spaced from the MCMC run) for each of the two independent runs, yielding a total of 10 replicates. For all simulations, the full model (with *K=4* conditions) was then applied to these simulated data. For a given pair of condition (e.g. HLAB57+ versus HLAB57-), and for several levels *α*, the number of positions inferred to be under differential selection with posterior probability greater than *1-α* was determined. In the context of the first series of simulations (no differential selection simulated), dividing this number by the total number of positions times the number of amino acids gives the rate of false positives, which was tabulated for several values of *α*. For the second series of simulations (with differential selection simulated), the discoveries made at a given threshold were compared with the true differential selection values, and the rate of false discovery was thus determined and plotted as a function of the significance threshold.

## Results

### Simulation analyses

The properties of the model were first investigated through simulations. Since the main application of the model introduced here is to identify positions for which specific amino acids are under condition-dependent selection pressure, the simulation analyses were more specifically designed to evaluate the power of this selection method, as well as its rates of false positives and of false discoveries. In order to ensure that the conclusions of the simulations are relevant for the empirical situations considered here, simulations were calibrated against parameter estimates obtained from the empirical analyses on the HIV dataset. This was done using the posterior predictive formalism (see methods). A first series of 10 simulations were conducted under the null model assuming no differential selection effect across conditions — thus, assuming a constant fitness landscape over the whole phylogenetic tree. The model with *K=4* conditions (see methods) was then applied to these simulated data. For a given pair of condition (e.g. HLAB57+ versus HLAB57-), and for different levels α, the number of positions inferred to be under differential selection with posterior probability greater than *1-α* was determined, giving us an estimate of the rate of false positives as a function of the stringency of the selection. As can be seen from table 1, for reasonable posterior probability thresholds, the rate of false positive is low, reaching 5% for *1-α* = 0.7, and virtually equal to 0 for *1-α* > 0.8.

**Table 1.** Rate of False Positive for different conditions and different thresholds.

| **threshold interval** | **Within-patient** | **B57<sup>+</sup> patients** | **B35<sup>+</sup> patients** |
| --- | --- | --- | --- |
| **0.5 - 0.55** | 186.94 | 194.37 | 192.82 |
| **0.55 - 0.6** | 22.28 | 10.14 | 12.76 |
| **0.6 - 0.65** | 6.62 | 2.05 | 2.88 |
| **0.65 - 0.7** | 2.01 | 0.36 | 0.60 |
| **0.7 - 0.75** | 0.26 | 0.05 | 0.06 |
| **0.75 - 0.8** | 0 | 0 | 0 |
| **0.8 - 0.85** | 0 | 0 | 0 |
| **0.85 - 0.9** | 0 | 0 | 0 |
| **0.9 - 0.95** | 0 | 0 | 0 |
| **0.95 - 1** | 0 | 0 | 0 |

This simulation experiment illustrates an important point about the Bayesian approach used here: the use of a normal distribution centered on 0 enforces shrinkage of the differential fitness effects across positions towards 0 (i.e. the model is centered on the null hypothesis representing an absence of selective difference between conditions). One important consequence of this choice is that, in the absence of a sufficiently strong empirical signal able to counteract this prior, the method will typically not infer high posterior probability support for differential selective effects.

A second series of simulations was conducted, under the full model, i.e. assuming the presence of modulations of the fitness landscape across conditions. The true values of the differential selection effects defined by these simulations were set aside and the 4- condition model was then applied to each of the 10 simulation replicates. For a given pair of condition (e.g. HLAB57+ versus HLAB57-), and for a given level *α*, the set of discoveries at level *α* (i.e. the set of all positions/amino acid pairs such that the posterior probability of a differential selection effect between the two conditions is greater than *1-α*) was determined. A discovery was then deemed to be false if the true selective effect for that amino acid at that position is of the opposite sign as the one inferred by the model. The rate of false discovery (FDR) was plotted as a function of *1-α* in figure 2. As expected, the FDR decreases with the stringency of the test. For model M1 at threshold around 0.80, the FDR lies around 15%. For a threshold of 0.90, the FDR is around 10% and reaches about 5% for the B57+/B57− comparison. For model M2, the FDR value is higher for the B57+/B57− and B35+/B35−, compared to M1. In within-patient condition, the two models produce very similar FDR values. Based on these simulations, in the following, we use a two-level selection procedure, with two thresholds at 0.80 and 0.90. We will refer to the corresponding discoveries as moderately and strongly supported findings, respectively.

**Figure 2.**
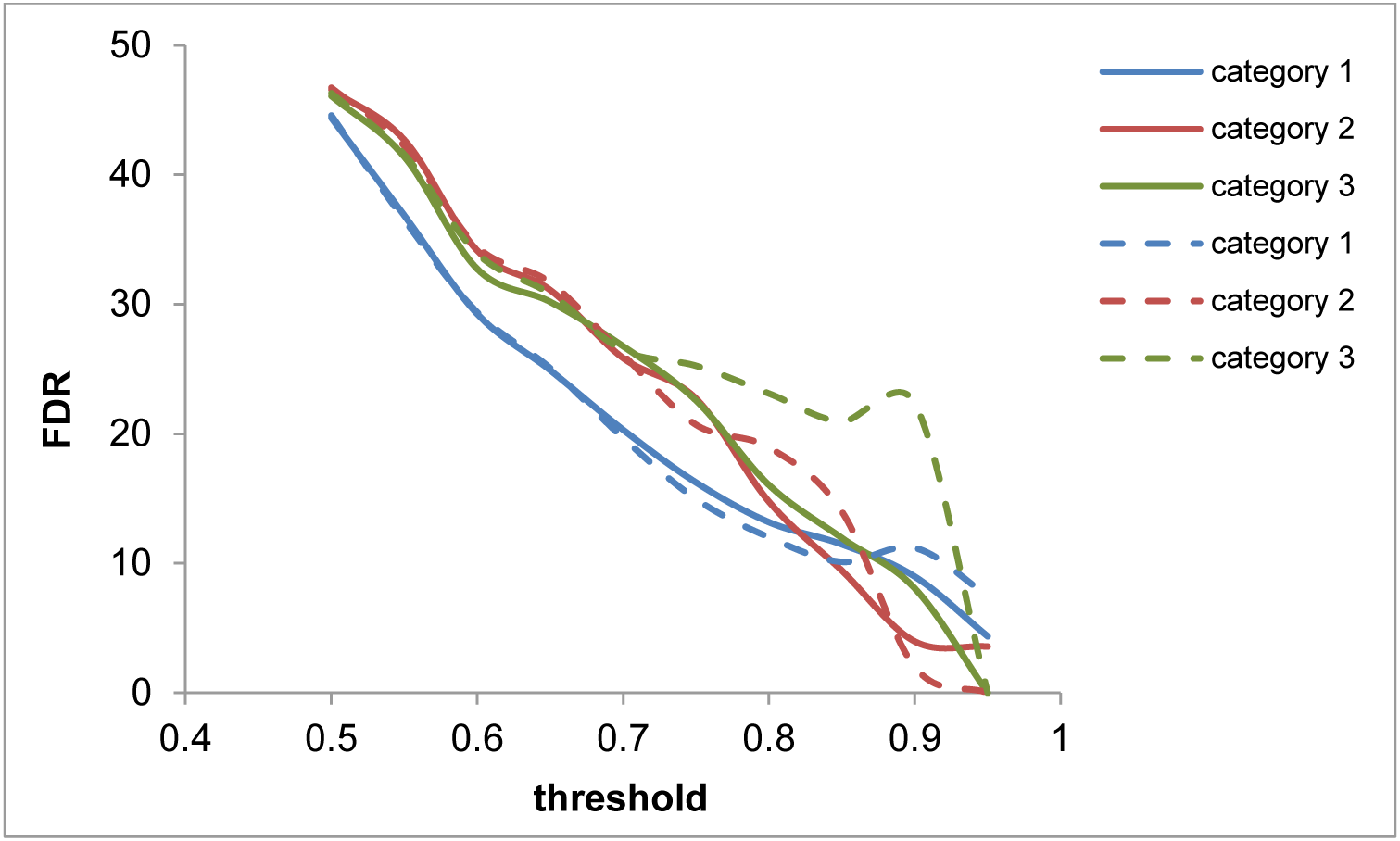
FDR according to posterior probability threshold for 3 categories for model M1 (line) and M2 (dash line). category 1 (blue), 2 (red) and 3 (green) represent within, B57+ and B35+ patients.

### Analyses of HIV empirical data

Our DS model was applied to a dataset of HIV coding sequences (encoding the Gag protein) obtained from 41 patients (see Methods). This dataset is interesting for two reasons. First, it contains multiple sequences (8 on average) for each patient, thus providing empirical information about within-host evolution of viral genetic sequences. Second, the HLA types of the patients is known, and therefore, it is possible to correlate the amino acid patterns observed in viral sequences with the HLA type of the host.

The evolution of HIV-1 is characterized by a complex interplay between short-term and long-term molecular evolutionary processes. Short-term evolution takes place mostly within hosts. It involves a selection pressure for fast replication and for efficient escape from the immune system. Long-term evolution, on the other hand, involves repeated switches between hosts. As a result, the selective forces involved in long-term evolution depend not only on replication but also on the infectivity of the virus and on its ability to adapt to a constantly changing immunological environment. Short- and long-term evolution are also characterized by different population-genetic regimes: within-host populations contain a substantial fraction of segregating polymorphism, some of which are potentially deleterious and therefore ultimately eliminated by purifying selection, whereas most differences having occurred along the branches connecting host-specific groups of sequences have essentially reached fixation and are therefore probably either nearly-neutral or adaptive. Of note, there is a bottleneck occurring as the disease transmits from one individual to another (between-patient). This bottleneck at transmission, which has been shown by the homogeneity of HIV-1 in very early infection [32-34], probably contributes to the reduction of segregating polymorphisms between-hosts molecular evolution. Altogether, the interplay between these two evolutionary timescales results in a complex process, potentially involving selective conflicts between short-term within-host competition and long-term survival in an immunologically highly polymorphic human population.

Accordingly, in this study, we partitioned the phylogenetic tree relating the viral sequences into different categories: first, we distinguished between the branches connecting the host-specific groups of sequences (between-patient condition) and the branches within each host-specific group of sequences (within-patient condition). Among the latter set of branches, we further distinguished among patients according to their HLA-type: either between HLA-B57+ and HLA-B57-patients, or between HLA-B35+ and HLA-B35-patients. The HLA-B57 type is known to be associated with the control of viremia [35, 36] whereas HLA-B35 is known as the HLA related to the fast progression of the disease [37, 38].

A global reference selection profile is estimated by our method. This reference fitness landscapes, which captures the baseline site-specific amino acid preferences in the form of site-specific vectors of 20 fitness factors (one for each amino acid), can be visualized using a graphical logo representation [39] and compared with the reference HIV-1 sequence (HXB2, the first 60 coding positions are shown in Figure 3). The selection profile inferred with our method is highly similar to the reference sequence (the fittest amino acid corresponds to the amino acid of the reference sequence at 86% of the coding positions). In some cases, compared to the reference sequence, the fitness profile suggests a distinct but biochemically similar dominant amino acid (e.g. position 15, K instead of R), or several equally fit amino acids (e.g. position 30). This corresponds to the actual sequence variation observed in our empirical alignment. Altogether, this global reference selection profile illustrates that HIV evolution occurs on a background characterized by strong purifying selection, allowing for a very limited set of amino acid sequences for the viral protein.

**Figure 3.**
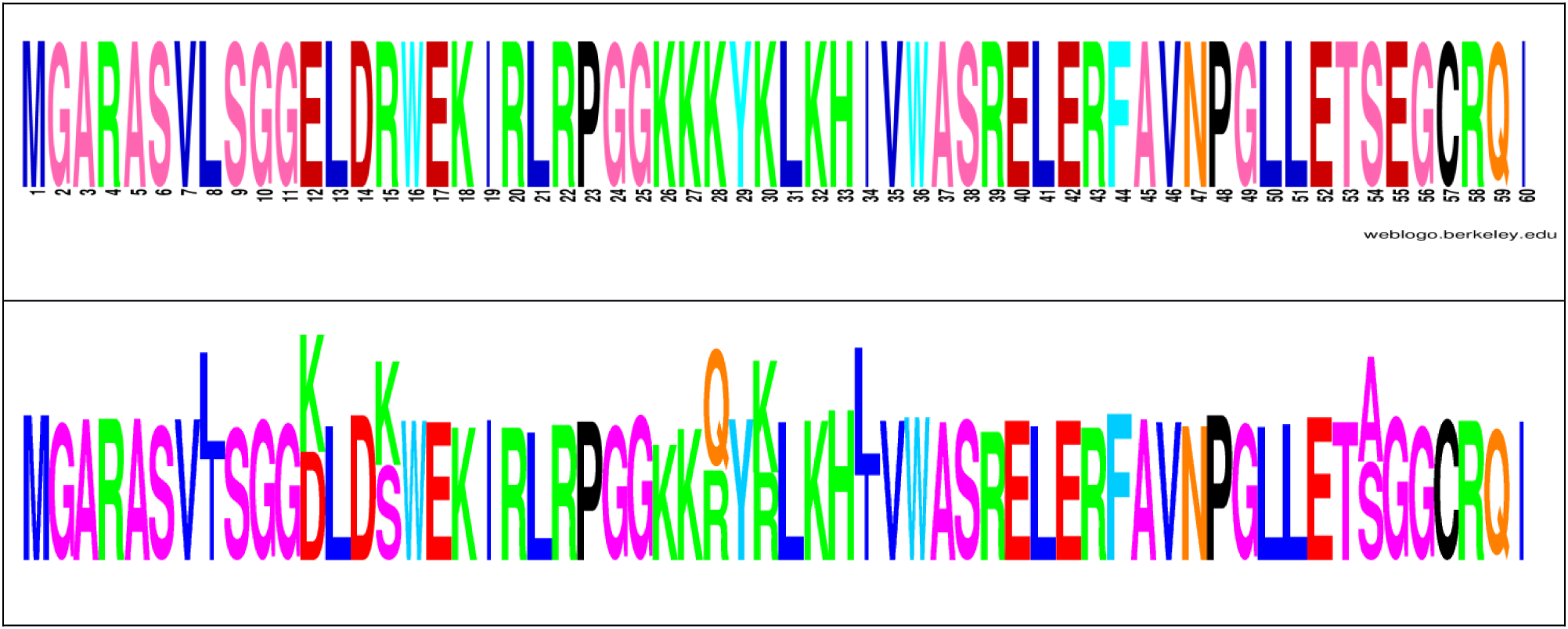
Comparison of HIV-1 global selection profile estimated by DS model with the reference sequence HXB2. The first 60 amino acids are shown. HXB2 sequence is at the top and global selection profile is at the bottom. The reference logo was made using Weblogo [40].

Against this background fitness landscape, our model then estimates differential selection profiles between each pair of conditions: first, between within-host and between-host (Figure *4-b* and *5-b*), and second, among within-host sequences, between HLA-B57- and HLA-B57+ sequences (Figure *4-c*), or between HLA-B35- and HLA-B35+ sequences (Figure *5-c*). The logos represented on Figure 4 and 5 indicate whether the fitness of any particular amino acid is inferred to be increased (above the line) or decreased (below the line), at a given position, between the two conditions being compared. These figures only give point estimates for the differential effects. In practice, the posterior probability support associated to these estimates is most often relatively low (figure 6), except for a small subset of positions for which stronger evidence (p.p. > 0.8) for a differential selection effect is inferred by the model. These more clear-cut cases represent our findings, which are given in Table 4 for the two model settings.

**Figure 4.**
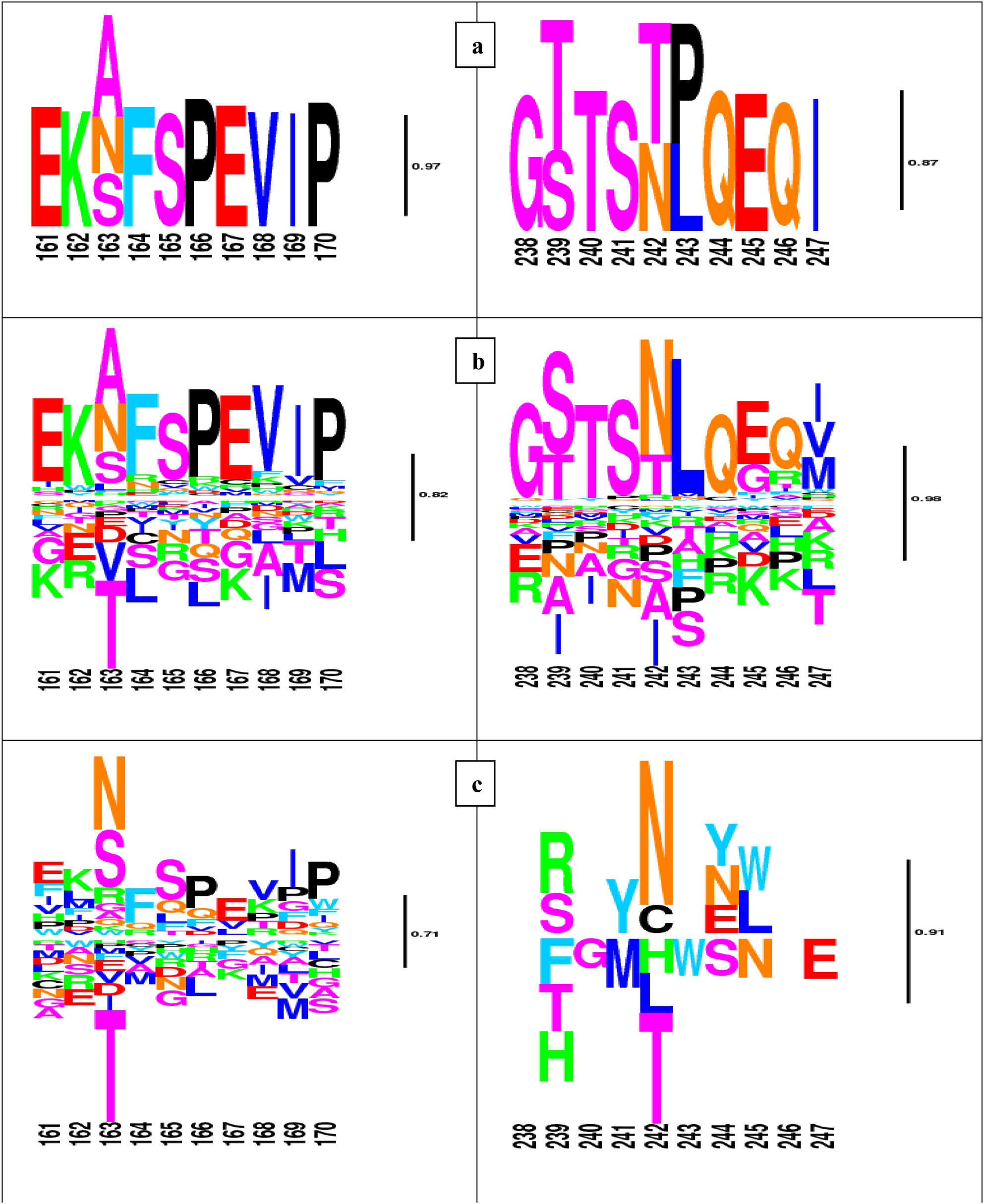
Global and differential selection profiles (differential for HLA-B57). a. Global selection profile (G). b and c. Differential selection profile for within-patient and HLA-B57+ group, respectively. The posterior probability of positive selection for N and negative selection for T at position 242 (TW10 epitope) is 0.94 and 0.88 in HLA-B57+ hosts. At position 163 (KF11 epitope), N is selected positively with the posterior probability of 0.77. The logos are filtered for p.p. below 0.05. Heights are proportional to posterior mean differential selective effects.

**Figure 5.**
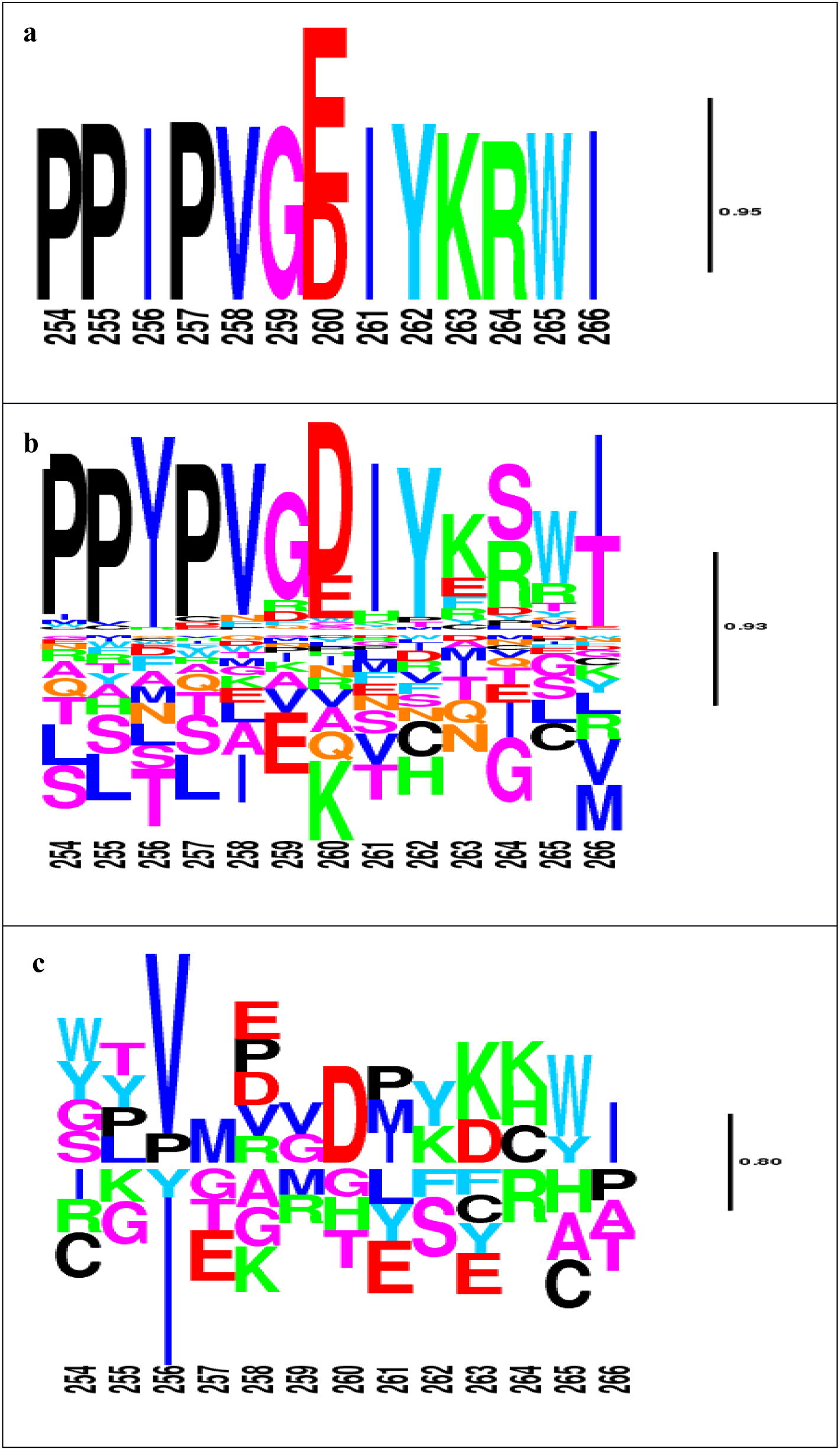
Global and differential selection profiles (differential for HLA-B35). **a.** Global selection profile of NY10 epitope (251-260). The epitope’s global selection is 100% matching the HXB2 sequence. **b** and **c.** Differential selection for within-patient HLA-B35- and HLA-B35+, respectively. The posterior probability of E to D substitution at position 260 is 0.81. The logos are filtered for p.p. less than 0.05. Heights are proportional to posterior mean differential selective effects.

**Figure 6.**
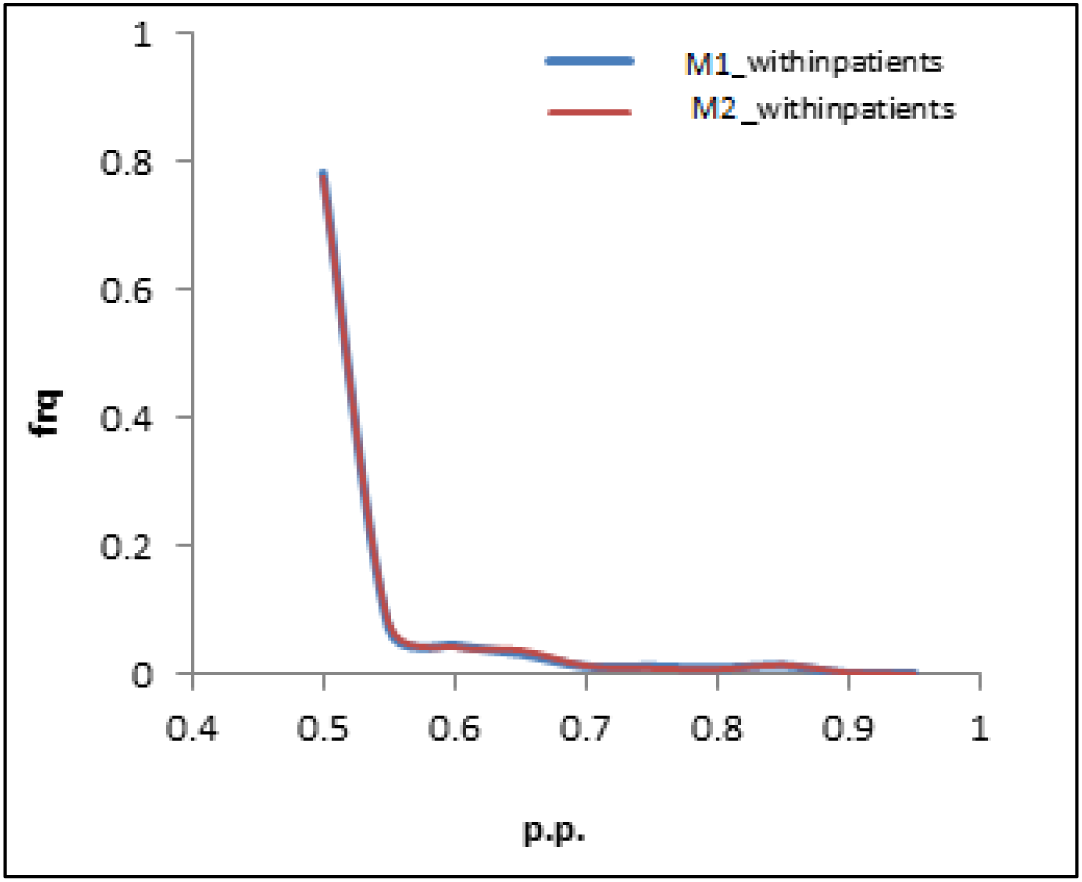
Frequency plot for posterior probabilities of differential selection effects across all amino acids at all positions; phenomenological (M1) versus mechanistic (M2) approach.

From Table 4, we see that, by far, the largest number of differentially selected amino acid variants is found when comparing the within- and between-patient conditions, with more than 280 findings under both models. On the other hand, a quick look at the corresponding profiles suggests that this is mostly due to a global difference in the intensity of selection (or a global difference in statistical power), rather than to specific selective differences between the two conditions (see discussion).

The differences between alternative HLA backgrounds, on the contrary, seem to be more specific. These findings are listed with more details (position, amino acid, credible interval and posterior probability support) in Table 2 and Table 3 for B57+ and B35+ conditions, respectively. Among them, there are some known mutations identified in association with specific HLAs. Two important HIV-1 escape mutations defined in B57+ patients are T242N and A163X in epitopes TW10 [41, 42] and KF11 [43, 44], respectively. X at position 163 is mostly P and N. The logos of the corresponding regions are shown in Figure 4. The selection factors estimated at these positions are in agreement with these previously known escape mutations.

**Table 2.** List of differentially selected amino acids for B57+ hosts with p.p.> 0.80.

| position | amino acid | p.p. | median | lower | upper | fitness |
| --- | --- | --- | --- | --- | --- | --- |
| 242 | N | 0.93 | 1.36 | -0.37 | 3.07 | increased |
| 248 | G | 0.91 | -1.20 | -2.82 | 0.45 | decreased |
| 30 | Q | 0.89 | 1.09 | -0.69 | 2.92 | increased |
| 242 | T | 0.87 | -0.95 | -2.55 | 0.78 | decreased |
| 30 | K | 0.87 | -0.96 | -2.49 | 0.69 | decreased |
| 357 | A | 0.86 | 0.94 | -0.73 | 2.86 | increased |
| 15 | R | 0.86 | 0.72 | -1.01 | 2.41 | increased |
| 118 | A | 0.85 | -0.93 | -2.69 | 0.79 | decreased |
| 239 | S | 0.85 | 1.02 | -0.95 | 2.64 | increased |
| 137 | L | 0.82 | -0.86 | -2.55 | 0.93 | decreased |
| 326 | S | 0.81 | 0.79 | -1.28 | 2.46 | increased |
| 357 | G | 0.81 | -0.78 | -2.55 | 0.97 | decreased |
| 280 | T | 0.80 | 0.83 | -0.79 | 2.43 | increased |
| 12 | E | 0.80 | 0.71 | -0.96 | 2.43 | increased |
| 248 | E | 0.80 | 0.66 | -0.97 | 2.42 | increased |
| 223 | I | 0.80 | -0.70 | -2.28 | 1.02 | decreased |

**Table 3.** List of differentially selected amino acids for B35+ individual with p.p.> 0.80.

| position | amino acid | p.p. | median | lower | upper | fitness |
| --- | --- | --- | --- | --- | --- | --- |
| **46** | L | 0.97 | 1.69 | -0.05 | 3.44 | increased |
| **34** | L | 0.96 | 1.52 | -0.31 | 3.19 | increased |
| **252** | H | 0.96 | 1.59 | -0.18 | 3.28 | increased |
| **111** | S | 0.93 | -1.15 | -2.72 | 0.49 | decreased |
| **127** | Q | 0.93 | -1.11 | -2.74 | 0.48 | decreased |
| **376** | V | 0.93 | 1.16 | -0.49 | 2.68 | increased |
| **312** | D | 0.92 | 1.23 | -0.55 | 3.06 | increased |
| **137** | M | 0.92 | 1.26 | -0.47 | 3.22 | increased |
| **252** | N | 0.92 | -1.05 | -2.60 | 0.48 | decreased |
| **30** | K | 0.92 | -1.05 | -2.44 | 0.52 | decreased |
| **248** | A | 0.91 | 1.25 | -0.41 | 3.07 | increased |
| **310** | T | 0.91 | 1.25 | -0.54 | 2.97 | increased |
| **441** | H | 0.89 | 0.95 | -0.43 | 2.46 | increased |
| **46** | V | 0.89 | -1.06 | -2.74 | 0.52 | decreased |
| **67** | A | 0.89 | 1.09 | -0.66 | 2.82 | increased |
| **111** | C | 0.88 | 1.08 | -0.75 | 2.76 | increased |
| **375** | V | 0.88 | -0.85 | -2.48 | 0.72 | decreased |
| **255** | V | 0.88 | 1.08 | -0.79 | 2.61 | increased |
| **441** | Y | 0.87 | -0.92 | -2.37 | 0.53 | decreased |
| **405** | I | 0.86 | 0.94 | -0.72 | 2.51 | increased |
| **15** | Q | 0.86 | 0.94 | -0.77 | 2.84 | increased |
| **138** | L | 0.86 | -0.90 | -2.41 | 0.76 | decreased |
| **376** | I | 0.85 | -0.81 | -2.26 | 0.67 | decreased |
| **127** | T | 0.85 | 1.01 | -0.86 | 2.83 | increased |
| **69** | Q | 0.84 | -0.79 | -2.37 | 0.78 | decreased |
| **81** | A | 0.84 | 0.94 | -0.74 | 2.65 | increased |
| **176** | A | 0.84 | 0.86 | -0.88 | 2.86 | increased |
| **280** | T | 0.83 | 0.96 | -0.86 | 2.40 | increased |
| **348** | S | 0.83 | 0.97 | -0.90 | 2.87 | increased |
| **61** | I | 0.83 | 0.77 | -1.14 | 2.61 | increased |
| **81** | T | 0.83 | -0.82 | -2.41 | 0.85 | decreased |
| **268** | M | 0.82 | 0.81 | -0.81 | 2.45 | increased |
| **280** | A | 0.82 | -0.82 | -2.41 | 0.85 | decreased |
| **388** | K | 0.82 | 0.74 | -0.90 | 2.37 | increased |
| **389** | P | 0.82 | 0.81 | -0.81 | 2.45 | increased |
| **397** | R | 0.82 | 0.72 | -1.00 | 2.53 | increased |
| **95** | R | 0.82 | 0.77 | -0.83 | 2.39 | increased |
| **68** | I | 0.81 | 0.87 | -1.14 | 2.67 | increased |
| **215** | L | 0.81 | -0.73 | -2.19 | 0.70 | decreased |
| **118** | T | 0.81 | 0.70 | -0.95 | 2.33 | increased |
| **260** | D | 0.81 | 0.75 | -1.00 | 2.48 | increased |
| **54** | A | 0.81 | 0.75 | -0.96 | 2.52 | increased |
| **93** | A | 0.80 | 0.73 | -1.04 | 2.44 | increased |
| **28** | K | 0.80 | -0.66 | -2.46 | 1.06 | decreased |
| **58** | K | 0.80 | 0.69 | -1.31 | 2.34 | increased |

**Table 4.** Numbers of differentially selected amino acid-positions with posterior probability >0.70 and >0.90 in different conditions estimated by models M1 and M2.

| Threshold | Model | Within-patient | B57<sup>+</sup> patients | B35<sup>+</sup> patients |
| --- | --- | --- | --- | --- |
| >0.80 | M1 | 281 | 15 | 48 |
| >0.80 | M2 | 286 | 5 | 30 |
| >0.90 | M1 | 54 | 2 | 13 |
| >0.90 | M2 | 56 | 0 | 1 |

Intriguingly, the T/N escape variant at position 242 (TW10 epitope) is not recovered by the mechanistic model (M2), suggesting that the phenomenological model is more adequate to predict differential selection patterns. This confirms our simulation studies, suggesting that the phenomenological model has a greater detection power. Also of interest, our method does not infer that T is preferred in a B57- environment, whereas N is favored in a B57+ background. Instead, it suggests that both amino acids are acceptable in a B57- environment, but that N becomes the only one favored in B57+ patients. A similar pattern is observed for the A163X escape mutation, with p.p. = 0.7. One known mutation for B35+ individuals is E260D in NY10 epitope [45]. Our method detects this mutation to be under condition-specific selection with posterior probability of 0.81 (Figure 5).

### Sensitivity to tree topology

As mentioned in the phylogenetic tree estimation section, we used three monophyletic topologies of the tree in the analysis. We refer to these trees as tree T1, T2 and T3. Having used them in M1-DS model, we found that the number and the key differentially selected positions are very similar for all trees. The number of these differentially positions is summarized for B57+ and B35+ patients and at each significance threshold in Table 5.

**Table 5.**
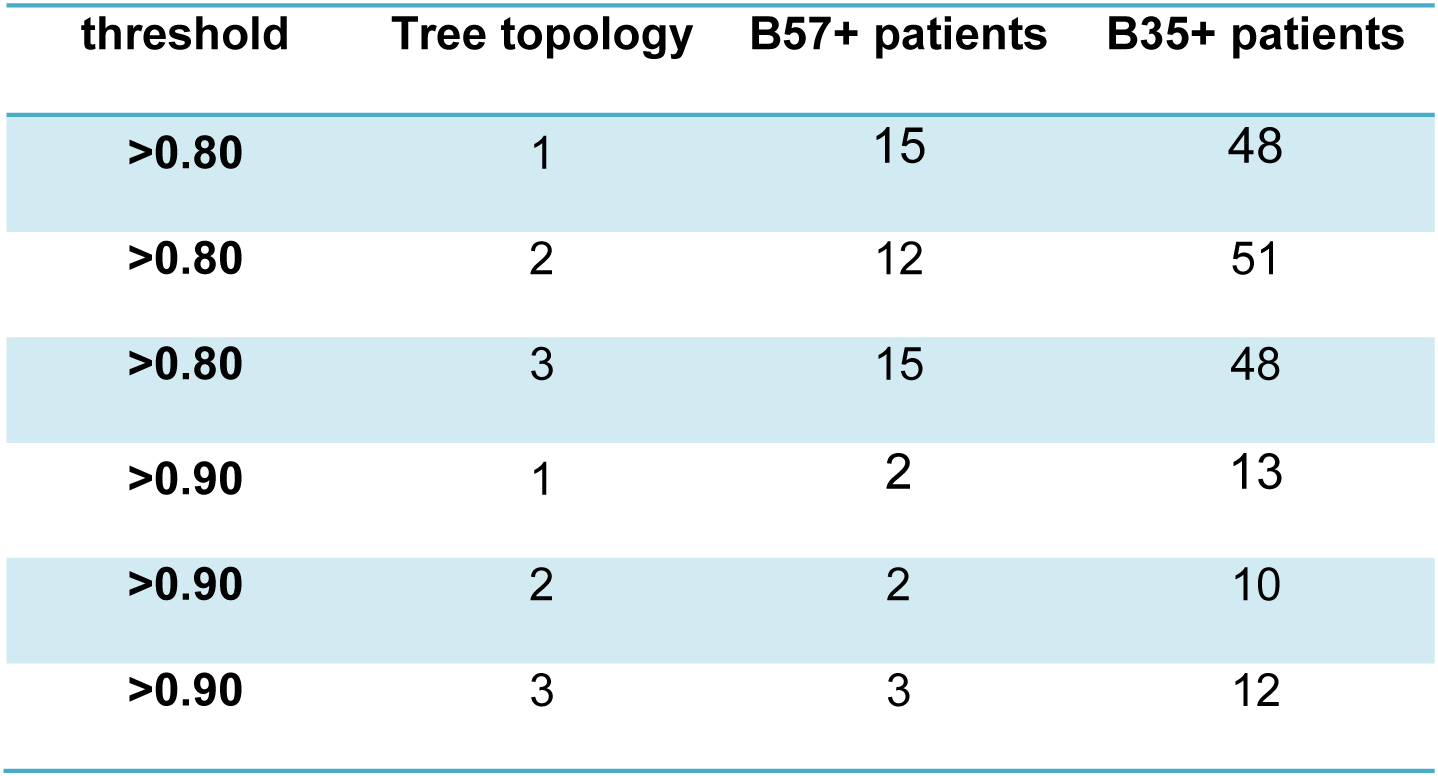
Number of differentially selected positions with posterior probability >0.80 and >0.90 obtained by M1-DS model using tree T2 and tree T3.

By comparing the positions declared significant for each threshold, we see that in B57+ condition, all findings under tree T1 (nj topology) were recovered under tree T2 (MrBayes topology with constraint) and tree T3 (MrBayes topology without constraint) for threshold greater than 0.85. Only 3 and 5 positions were not found at the threshold of 0.80 for trees T2 and T3, respectively. None of the positions found different between two topologies belong to the positions previously known to correspond to viral escape mutants.

Altogether, the relatively small number of sequences that had to be removed, combined with the relative robustness of our result to the exact choice of the tree topology, suggests that the problems of multiple infection patterns, or tree reconstruction errors, have a globally marginal impact on our analysis.

## Discussion

Here, we have introduced a hierarchical Bayes method for detecting adaptive patterns in protein coding sequences as a function of known selective backgrounds. Compared with previously introduced methods [21, 22], our approach has several additional features. The approach of Carlson et al [21], relying on a Bayesian network representation, is formulated at the codon level. In addition, it can accommodate epistatic effects (see below). Besides, it is focused on the terminal branches of the phylogeny and therefore ignores potentially relevant empirical information from the deeper parts of the phylogenetic tree. The approach of Tamuri et al [22], in contrast, fully integrates the empirical signal over the entire tree. However, it is formulated directly at the amino acid level and does not explicitly account for the coding structure. Our method has the strengths from these two approaches: like Carlson et al [21], it is formulated at the codon level; as in [22], it relies on an explicit evolutionary model with site- and condition-specific selective effects.

The fact that our method integrates the empirical signal about more ancient codon substitutions opens new possibilities, in particular, for comparing short-term (within-host) and long-term (between-host) adaptive patterns. As it stands, however, the results obtained in this direction are not yet so convincing: the within-host differential selection profiles obtained through our method (figures *4-b* and *5-b*) seem to partially reproduce the condition-independent amino acid fitness profiles (figures *4-a* and *5-a*). The reasons for such a redundant output are not totally clear. Deleterious mutations segregating within-host, but purified away in the long-term (and therefore absent from the deeper branches of the phylogeny connecting host-specific clusters) are an important difference between within- and between-host conditions. However, such segregating polymorphisms would be expected to result in an opposite pattern, leading to artefactual high selection coefficients in the within-host condition for unfit amino acids that are not observed in the between-host selection profiles. One alternative explanation for the observed redundancy would be that the law of condition-independent selection profiles across sites is not correctly captured by a Dirichlet distribution. Possibly for that reason, the remaining part of the condition-independent selective effects may be captured by the differential selection profile of the within-host condition. Ultimately, more sophisticated hierarchical Bayesian settings could be used, such as non-parametric priors [6]. The combination of condition- and site-specific effects is computationally challenging, and further algorithmic work is therefore needed in this direction to fully accommodate arbitrary distributions of random-effects across positions and conditions.

The distribution of differential selective effects across sites and conditions may also need additional statistical and computational developments in the long term. Here, we have used Normal distributions centered on 0 to model differential selective effects. Doing this leads to efficient soft shrinkage toward 0. However, this approach does not implement sparsity: All amino acids, at all positions and under all conditions, have non-zero differential selective effects with a posterior probability of one. Ultimately, sparse differential selection profiles (with only a small number of positions and amino acids displaying significant non-null differential selective effects with high posterior probability) could be obtained through the use a spike-and-slab mixture model [46].

Two alternative models of the rate of change between codons were considered in this study: one purely phenomenological [6, 9], and another one that has a better mechanistic justification, based on first principles of population genetics. When applied to HIV sequences, however, the mechanistic model does not seem to lead to better results, compared to the phenomenological approach. In particular, it fails at detecting known HLA-restricted escape mutations. The mechanistic model, however, makes several assumptions that are clearly not warranted in the present context: low-mutation approximation, and more fundamentally, a mutation-fixation paradigm [7, 47], which amounts to ignore clonal interference. In sharp contrast, viral sequences evolve under a very high mutation rate, leading to strong clonal interference. Another consequence of the very high mutation rate is that segregating deleterious polymorphisms are expected to be present at a substantial frequency, something which is not correctly captured by the mutation-selection model: fundamentally, this model is meant to be applied to inter-specific data. Here in contrast, a meta-population model would be more adequate. The theoretical and computational developments in this direction still appear to be challenging.

Our method does not take into account epistatic interactions between positions. Yet, those interactions seem to play an important role in HIV evolution, in particular concerning escape mutations. Most escape mutations cause a viral fitness cost which leads to decreased replication of the virus [41]. Position 242 is under the strongest selection pressure from the immune system which corresponds to the ability of B57+ hosts to control the disease. T242N mutation in B57+ individuals reverts in viruses transmitted to a HLA-mismatched host [42], which supports the fact that the mutation has a strong fitness cost for the virus in terms of replication capacity [48]. This fitness cost might be compensated for, to some extent, by mutations at other positions, mostly around the escape mutation. In sequences with T242N mutation, the compensatory mutations H219Q, I223V, M228I/V, G248A and N252H were identified [41, 42]. It has been reported that these mutations are significantly more frequent in HLA-B57+ patients with a progressing disease compare to HLA-B57+ non-progressors [41]. In this study, we did not see significant differences for final amino acids (Q, V, I/V, A and H) between B57+ and B57− patients at those suppressing positions, although initial amino acids are significantly unfavored (p.p.=0.80, 0.91, 0.77 for I, G and N at positions 223, 248 and 252, respectively). There may be two reasons for that; first, our model takes each site into account independently and codon co-variation is not considered. Secondly, contrary to escape mutations which revert in the HLA mismatch host, compensatory mutations do not tend to revert after transmission to HLA mismatch individuals [42]. For example, H219Q, the associated mutation to T242N, is reported to be maintained after transmission from B57+ to B57− hosts. So, this mutation might be stable and spread in the population. As it stands, explicitly implementing epistatic effects in the context of the present modeling framework appears to be challenging, although not impossible [49].

## Conclusion

We proposed a phylogenetic differential selection model, which is able to find adaptive patterns in coding sequences influenced by selective environments. Applying the model on HIV-1 *Gag* sequences, leads to the detection of a few amino acid-positions that are differentially selected under different host HLA types, as HIV tries to escape from immune system through its fast evolution. The model is thus able to find known HLA-restricted mutations, as well as some new mutations, to be under differential selection. The power of our model is that it is capable of detecting both positive and negative selection pressure on each amino acid at each position under each environmental condition.

This differential selection model can be used in other situations in which differential selective effects are suspected, as a function of known predictors, for viruses (e.g. finding adaptive patterns of HIV sequences under the selection pressure of immune system or antiviral therapy provides an insight of the direction of HIV-1 evolution in different hosts with different genetic characteristics), or in other species (e.g. convergent adaptations of multiple lineages of plants, or animals, to specific environmental conditions [50].

## Abbreviations

DS: Differential selection
HLA: Human leukocyte antigen
CTL: Cytotoxic T lymphocyte
MCMC: Markov Chain Monte Carlo
LANL: Los Alamos National Laboratory

## Authors’ Contributions

SP and NL conceived the project and participated in its design. SP performed the experiments. SP and NL analyzed the results. SP drafted and NL edited the manuscript. Both authors read and approved the final manuscript.

## Acknowledgement

We are thankful to the Natural Sciences and Engineering Research Council of Canada (NSERC) for financially supporting this research. We also thank the anonymous reviewers for their comments on the manuscript.

